# Nitroimidazopyrazinones with oral activity against tuberculosis and Chagas disease in mouse models of infection

**DOI:** 10.1101/2022.04.05.487095

**Authors:** Chee Wei Ang, Brendon M. Lee, Colin J. Jackson, Yuehong Wang, Scott G. Franzblau, Amanda F. Francisco, John M. Kelly, Paul V. Bernhardt, Lendl Tan, Nicholas P. West, Melissa L. Sykes, Alexandra O. Hinton, Raghu Bolisetti, Vicky M. Avery, Matthew A. Cooper, Mark A.T. Blaskovich

## Abstract

Tuberculosis remains one of the leading causes of death from a single infectious agent, surpassing both AIDS and malaria. In recent years, two bicyclic nitroimidazole drugs, delamanid and pretomanid have been approved to treat this airborne infection. This has spurred a renewed interest in developing new and improved nitroimidazole analogs. We have previously identified a new bicyclic heteroaromatic subclass, the nitroimidazopyrazinones, with substituted analogs showing promising activity against *Mycobacterium tuberculosis* under both aerobic and hypoxic environments. A second generation of nitroimidazopyrazinones with extended biaryl side chain also possessed good antiparasitic activity against *Trypanosoma brucei brucei* and *Trypanosoma cruzi*, suggesting the utility of this new scaffold for development into potential candidates against both tuberculosis and the kinetoplastid parasites which cause neglected tropical diseases. In this study, we further evaluated the properties of nitroimidazopyrazinone derivatives by assessing their selectivity against different mycobacterial species, measuring their reduction potential, and determining the kinetic parameters as substrates of the deazaflavin-dependent nitroreductase (Ddn), which is the activating enzyme of delamanid and pretomanid in *M. tuberculosis*. We also conducted an *in vivo* evaluation of a lead compound, MCC8967 that demonstrated a favorable pharmacokinetic profile, with good oral bioavailability and efficacy in an acute *M. tuberculosis* infection model. Two other promising compounds MCC9481 and MCC9482, with good *in vitro* activity (IC_50_ = 0.016 and 0.10 µM, respectively) against *T. cruzi*, the causative agent for Chagas diseases, were similarly tested for *in vivo* activity. These compounds also exhibited good oral bioavailability, and transiently reduced the acute-stage parasite burden by >98‒99% at doses of 50 mg/kg once or twice daily, similar to benznidazole at 100 mg/kg once daily. Overall, we have demonstrated that active nitroimidazopyrazinones have potential to be developed as clinical candidates against both tuberculosis and Chagas disease.

**Author Summary:** Tuberculosis and parasitic infections continue to impose a significant threat to public health and economic growth worldwide. Most of the efforts to control these diseases still rely on drug treatments with limited effectiveness and significant side effects. There is now an urgent need to develop new treatments to combat these infections. Here, we report the *in vitro* and *in vivo* profile of a new bicyclic nitroimidazole subclass, namely nitroimidazopyrazinones, against mycobacteria and *Trypanosoma cruzi*. We found that derivatives with monocyclic side chains are selective against *Mycobacterium tuberculosis*, the causative agent of tuberculosis, but not active against other nontuberculosis mycobacteria. In an acute mouse model, they were able to reduce the bacterial load in lungs via oral administration. From a biochemistry perspective, we demonstrated that deazaflavin-dependent nitroreductase (Ddn) could act effectively on nitroimidazopyrazinones, indicating the potential of Ddn as an activating enzyme for these new compounds in *M. tuberculosis*. We also showed that derivatives with extended biaryl side chain were effective in suppressing infection in an acute *T. cruzi* infected murine model, with satisfactory oral bioavailability. These findings improve the understanding of the biological profile of nitroimidazopyrazinones for further development as potential antitubercular and antiparasitic agents.

## Introduction

Tuberculosis (TB) is one of the world’s deadliest infectious diseases. It is caused by *Mycobacterium tuberculosis*, a slow growing bacterial pathogen that was discovered by Dr. Robert Koch in 1882. According to the World Health Organization (WHO), it was estimated that 10 million people developed TB in 2020, with a death toll of 1.5 million [1]. In most cases, TB is treatable with a standard 6-month drug regimen comprised of isoniazid, rifampicin, ethambutol and pyrazinamide [1]. However, with the inexorable increase in drug resistance over the past decades, many first-line or even second-line antitubercular agents are no longer effective. It was reported that treatment success rates for rifampicin-resistant or multidrug-resistant TB were 55–59% in 2015-2018, compared to an average of 85% for drug-susceptible TB [1]. Following the theme of World TB Day 2021 - ‘The Clock is Ticking’, there is a pressing need to find new treatment options to end this old millennium disease. Given that current treatment regimens are long and complicated, which then lead to poor patient compliance, the desired target product profiles (TPPs) of new TB drugs focus on shortening the treatment duration, reducing the dosing frequency, enabling easy administration (ideally once a day oral dosing), and having the ability to overcome drug-resistant strains [2, 3].

Alongside TB and other more commonly known infectious diseases such as human immunodeficiency virus (HIV) and malaria, neglected tropical diseases (NTDs) are known as ‘infectious diseases of poverty’ as they affect the world’s poorest population [4]. Chagas diseases (CD), or American trypanosomiasis, affects 6‒7 million people worldwide and causes a huge annual economic loss of $7.2 billion [5, 6]. It is caused by the trypanosomatid parasite *Trypanosoma cruzi* that is transmitted mainly through a triatomine bug known as ‘kissing bug’ [7]. Although CD primarily affects Latin American countries, in recent years it has become a health concern for other continents including Europe, North America, and Australia due to population migration [6, 8]. Current therapeutic options for CD are limited to benznidazole and nifurtimox, drugs that were discovered in the 1960s and1970s. However, they suffer from various adverse effects, require long treatment duration, and have dubious efficacy in chronic infections [9]. Hence, ideal TPPs focus on new regimens that have a shorter treatment course, are safe and more efficacious than current drugs, exhibit good oral bioavailability, and are active against both chronic and acute phases [10]. In recent years, combination therapies that utilize multiple drugs with different mechanisms of action have been found to be effective in preclinical testing [11, 12]. Whilst combinations have been tested in clinical trials, their efficacy compared to individual drug components alone have yet to be identified [12].

There are many challenges in the area of TB and CD drug discovery. In the case of TB, hits identified through target-based screening almost always fail to show sufficient or comparable whole-cell activity. One of the reasons is due to the waxy, lipid-rich mycobacterial cell wall with low permeability, preventing most potential antibiotics from passing through and reaching the intracellular molecular targets [13]. In addition, efflux systems extrude a range of substrates, including drugs, out of the cells. In contrast, for *T. cruzi* there is lack of well-validated targets, thus most drug discovery efforts focus on phenotypic screening, providing cell permeable hits [14]. Fexinidazole was “re-discovered” by the Drugs for Neglected Diseases *initiative* (DND*i*) using phenotypic-based assays, and fexinidazole was ultimately evaluated as a clinical candidate for both human African trypanosomiasis (HAT) and CD [15, 16]. Two of the most recently approved TB drugs, delamanid and pretomanid, were also identified using phenotypic approaches [17]. They were later found to have repurposing potential against visceral leishmaniasis (VL) [18, 19], which is caused by the kinetoplastid protozoan, *Leishmania*. Through a collaborative program between the University of Auckland, TB Alliance, and DND*i*, a backup library of 900 pretomanid analogs have been counter-screened against *Trypanosoma brucei* and *T. cruzi,* resulting in the discovery of several promising hits to be pursued [20, 21].

Fexinidazole, delamanid, and pretomanid (Fig 1) all belong to the nitroimidazole family of drugs, which has a long history of use as treatments against bacterial and parasitic infections [22]. They are prodrugs that require bioactivation of the nitro group to exert their antimicrobial activity. The bicyclic nitroimidazole drugs, delamanid and pretomanid are active against both actively growing and non-replicating mycobacteria, making them an ideal candidate to shorten treatment duration [23]. They have interesting multiple modes of action. Under aerobic condition, these compounds target the mycobacterial cell wall by inhibiting the biosynthesis of mycolic acid [24]. When mycobacteria are in a hypoxic, non-replicating state, they cause respiratory poisoning by releasing nitric oxide (NO) intracellularly through bioactivation of the nitro group [25]. It has recently been found that a protein encoded Rv3547 is responsible for the reductive activation of delamanid and pretomanid in *M. tuberculosis* [26, 27]. This enzyme is now known as deazaflavin-dependent nitroreductase (Ddn) [28]. The Ddn mediated metabolism of pretomanid produced three stable metabolites, and one of the major products is a des-nitro species, which correlates well with the release of NO and its anaerobic bactericidal activity [25]. However, the Ddn enzyme is absent in kinetoplastids such as *T. cruzi* and *Leishmania*. It has recently been found that bicyclic nitroimidazoles are activated by a novel nitroreductase (NTR2) in *Leishmania* [29]. The NTR2 *Leishmania* enzyme could not activate the monocyclic nitroimidazole fexinidazole, which instead was mediated by a type I nitroreductase (NTR1) [30]. Although the metabolic fate of fexinidazole in *T. b. brucei* and *T. cruzi* remains unknown, diminished NTR1 activity resulted in resistance to fexinidazole and cross-resistance to other nitroaromatic drugs including nifurtimox [31]. This suggests that there are diverse modes of action of nitroimidazoles between different organisms.

**Fig 1.**
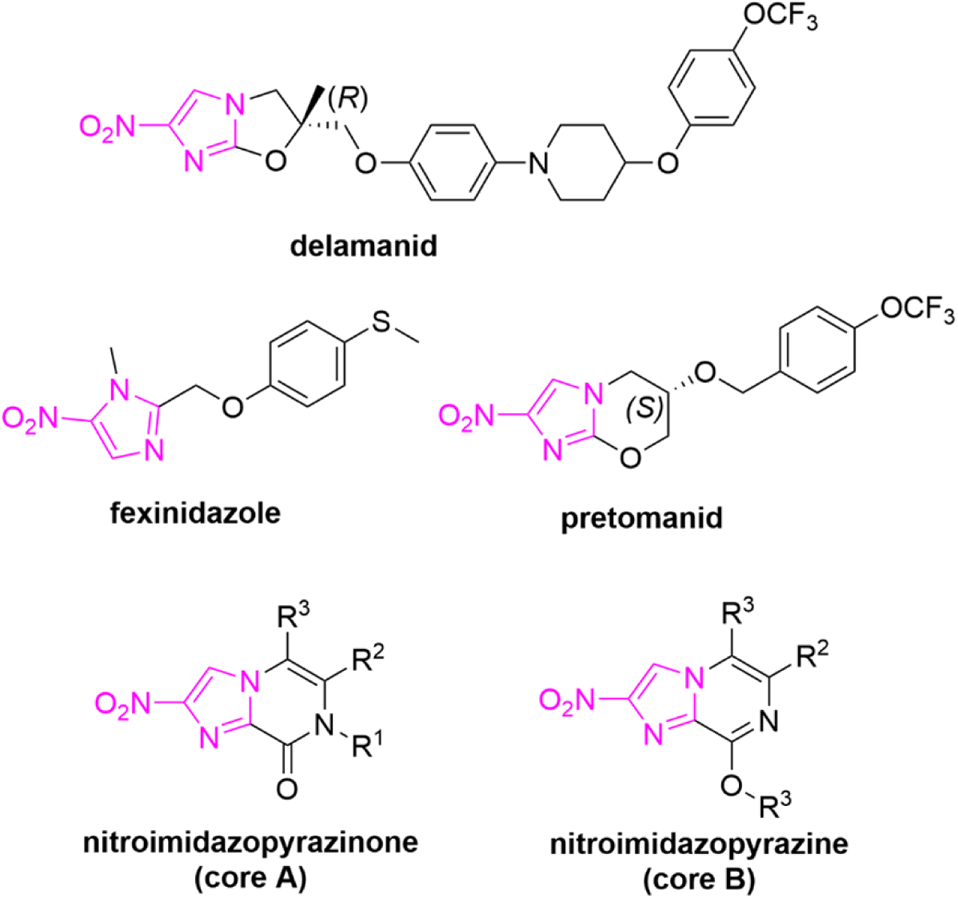
Chemical structures of monocyclic and bicyclic nitroimidazoles with antitubercular and antitrypanosomal activity.

We have previously described the synthesis of two novel regioisomeric bicyclic nitroimidazole scaffolds, the nitroimidazopyrazinones and the nitroimidazopyrazines (Fig 1). The nitroimidazopyrazinones in particular possessed antitubercular activity under both aerobic and hypoxic conditions (MIC = 0.06–32 μg/mL) [32]. Several promising hits were found to have comparable or better *in vitro* activity than pretomanid, while displaying low cytotoxicity against mammalian cell lines, good metabolic stability and high Caco-2 permeability. The *in vitro* activity profile of nitroimidazopyrazinones led to the current study, in which further evaluation of safety, efficacy and pharmacokinetic properties of this scaffold in murine models for both Chagas disease and tuberculosis have been undertaken. Our earlier investigations revealed the importance of the nitro group in the nitroimidazopyrazinones for antimicrobial activity, as the des-nitro analogs were as inactive as des-nitro pretomanid against *M. tuberculosis* [33]. Considering the correlation between the electrochemical reactions at the interface of electrode-electrolyte and the enzymatic redox reaction [34], we also now report the redox behavior of nitroimidazopyrazinones and their related subclasses using cyclic voltammetry (CV). To understand the possible modes of action of these new compounds against *M. tuberculosis*, we investigated whether they are activated by the same enzyme as pretomanid by conducting a biochemistry study using purified Ddn that is overexpressed in *Escherichia coli*. Lastly, we have explored the potential of selected nitroimidazopyrazinones in treating *T. cruzi* infection *in vivo*. Our previous *in vitro* results showed that the extended biaryl series could inhibit *T. cruzi* intracellular amastigotes, with IC_50_ values of ≤0.1 μM [35]. Here, we report the *in vivo* profile of two nitroimidazopyrazinones against *T. cruzi*, compared to the existing drug, benznidazole.

## Results and discussion

### Antimycobacterial activity of nitroimidazopyrazinones

In our previous studies, a library of 71 nitroimidazopyrazinones and 13 of their *O*-alkylated regioisomers (nitroimidazopyrazines) were synthesized and tested for their activity against *M. tuberculosis* H37Rv strain using a resazurin reduction assay [32, 35]. Given the encouraging activity demonstrated by the monocyclic side chain of nitroimidazopyrazinones under both normoxic and hypoxic conditions, additional *in vitro* susceptibility testing was performed to investigate their selectivity in different mycobacterial species (Table 1). Although *Mycobacterium smegmatis* has been used as a model system in most phenotypic screenings [36, 37] and its orthologs share 70% identity with *M. tuberculosis* genomes [38], our compounds displayed no activity against this non-pathogenic strain, highlighting the dangers of screening with a surrogate organism. They were also non-active against the opportunistic pathogen *Mycobacterium avium* when tested at concentrations up to 32 µg/mL under normoxic condition. These results were similar to pretomanid, suggesting the potential of these molecules as selective chemical starting points for development of therapeutics for diseases caused by *M. tuberculosis* complex but not against the nontuberculous mycobacteria. Previous studies have shown that Ddn orthologues, including those from *M. smegmatis* and *M. avium*, are inactive against pretomanid due to differences in sequence identity [39]. This could also explain the lack of activity of our nitroimidazopyrazinones in these mycobacteria when tested under normoxic condition.

**Table 1.** Minimum inhibitory concentrations (MICs) of nitroimidazopyrazinones against different mycobacterial species.

| Compound | Antibacterial<br>MIC (µg/mL)<br>normoxic |  |  |  | Antibacterial<br>MIC (µg/mL)<br>hypoxic |
| --- | --- | --- | --- | --- | --- |
|  | *M. tuberculosis*<br>H37Rv | *M. tuberculosis*<br>H37Ra | *M. avium* | *M. smegmatis* | *M. tuberculosis*<br>(H37Rv) |
| Pretomanid | 0.25-0.50 | ≤0.016-0.063 | >32 | >32 | 1 |
| MCC7270 | >32 | >32 | >32 | >32 | >32 |
| MCC7433 | 0.5 | 0.031-0.25 | >32 | >32 | 1-4 |
| MCC8364 | 0.13 | 0.13-0.25 | >32 | >32 | 0.5-2 |
| MCC8966 | 0.063 | ≤0.016 | >32 | >32 | 70-90% I @<br>0.125-8 µg/mL |
| MCC8967 | 0.063 | ≤0.016-0.031 | >32 | >32 | 0.25 |
| MCC8397 | >32 | 4-8 | >32 | >32 | >32 |
| MCC8399 | 1 | 0.5 | >32 | >32 | 4 |
| **MCC8400** | >32 | >32 | >32 | >32 | >32 |

Isoniazid control (normoxic): *Mtb* H37Rv MIC = 0.016 µg/mL, *Mtb* H37Ra MIC = 0.031-0.063 µg/mL, *M. avium* MIC = >32 µg/mL, *M. smegmatis* MIC = >32 µg/mL. Isoniazid control (hypoxic): *Mtb* H37Rv MIC = >5 µg/mL. Rifampicin control: *Mtb* H37Rv MIC = 0.005-0.01 µg/mL, *Mtb* H37Ra MIC = <0.016 µg/mL, *M. avium* MIC = 4 µg/mL, *M. smegmatis* MIC = 4 µg/mL. Rifampicin control (hypoxic): *Mtb* H37Rv MIC = 0.08-0.17 µg/mL.

To determine whether nitroimidazopyrazinones have similar activity against both virulent and avirulent organisms, we have compared their bioactivity against *M. tuberculosis* H37Rv and H37Ra strains under normoxic condition. Both strains are derived from the parent strain H37, but with some differences at their genomic and proteomic levels [40, 41]. It was reported that H37Ra acquired multiple mutations in genes that might be associated with its virulence attenuation [41, 42]. In our study, H37Ra was more susceptible to the tested analogs, with ~2–16-fold better activity compared to its virulent counterpart. Compounds MCC8966 and MCC8967 remained the most activity against both strains, with MICs of ≤0.016–0.031 µg/mL against H37Ra and 0.06 µg/mL against H37Rv under normoxic condition. When tested under low oxygen environment (0.1% oxygen), MCC8967 surpassed the activity of pretomanid against H37Rv (MCC8967: MIC = 0.25 µg/mL cf. pretomanid: MIC = 1 µg/mL). Activity against *M. tuberculosis* was only found for the pyrazinone scaffold, as the regioisomer nitroimidazopyrazine MCC8400 completely lost potency under both normoxic and hypoxic conditions. A similar structure-activity relationship (SAR) was observed between the two *M. tuberculosis* strains, where addition of substituents at R^2^ and R^3^ resulted in loss of activity against H37Rv and H37Ra strains. These results indicate that the less-hazardous H37Ra strain can potentially be used as a surrogate for the virulent *Mtb* H37Rv to screen for potential antitubercular hits, though most of the analogs tested generally demonstrated better activity against this avirulent model. However, this would need to be further validated under hypoxic condition, in addition to confirming with a broader range of compounds and drugs. As H37Ra can be handled in a biosafety level II environment, it would be advantageous to use this strain instead of *M. smegmatis* for primary screening, with promising hits then subjected to a more rigorous MIC determination using the pathogenic H37Rv strain, which requires level III containment.

### Electrochemical properties of nitroimidazopyrazin-one/-e scaffolds

We reasoned that the redox biochemistry of the nitro group is essential in mediating the biological activity of nitroimidazoles [43]. Pretomanid exhibits antitubercular activity through the loss of the nitro group, which in turn generates reactive nitrogen species within the mycobacterial cells to achieve anaerobic bactericidal activity [25]. However, the nitoimidazole redox potential should be optimum to exhibit high specificity against target organisms without causing unnecessary toxic effects to the hosts. For example, metronidazole possesses a more negative redox potential that exceeds the reduction capacity of mammalian redox systems and aerobic microbes, making it selective against anaerobes such as non-replicating, hypoxic *M. tuberculosis* [23]. 2-Nitroimidazoles, on the other hand, have a higher redox potential and are therefore more readily reduced by mammalian cells, making them more suitable as agents for cancer therapy [44]. One of the factors that affects the ease of nitro reduction is the electron affinity of the nitroheterocycles. As modification of molecular structure can affect the electrochemical properties and reduction potential of nitroimidazoles, the formal potentials (E’) of structurally related nitroimidazopyrazin-ones/-es were measured and compared using cyclic voltammetry (CV).

Initial measurements were undertaken in anhydrous DMSO as the reduction of nitroimidazoles is likely to be reversible under this condition, uncomplicated by coupled proton transfer. Representative cyclic voltammograms for metronidazole, pretomanid, and key compound MCC7433 are shown in Fig 2a. The E’ values were determined from the potential midway between the reduction and oxidation peaks, and calibrated against ferrocene. The peak-to-peak separation (> 59 mV and dependent on scan rate) was consistent with a quasi-reversible single electron transfer reaction to give the nitro radical anion as the product of electrochemical reduction. Metronidazole demonstrated a higher E’ than pretomanid (~200 mV difference, −1586 mV vs −1795 mV), which correlated well with a literature study [34]. The response for pretomanid was asymmetric (i_pc_ > i_pa_ at 200 mV/s) with an additional peak found upon re-oxidation at higher potential. At higher scan rates (> 1 V/s) the anodic and cathodic peak currents for pretomanid converged in magnitude, which is consistent with an irreversible chemical reaction following electron transfer. Investigations of related nitroimidazoles under similar conditions have suggested that irreversible dimerization of the radicals occurs [34]. Examination of the redox potentials of seven nitroimidazopyrazinones revealed a more positive E’ compared to pretomanid, with a range of −1555 to −1662 mV (Fig 2c, Table 2). MCC7433 displayed the highest potential (−1555 mV). The similarity of voltammograms exhibited by both metronidazole and MCC7433 indicated common electrochemical behavior. The addition of an electron-donating alkyl group at R^2^ and R^3^ (MCC8399) slightly increased the electron density of the nitroimidazole ring and consequently lowered its redox potential (−1566 mV). Pyrazine analog MCC8400 is a regioisomer of MCC8399 and showed an E’ value that was 30 mV more positive (−1536 mV). This also correlated well with the ^1^H NMR analysis, in which the methylene protons (‒O‒CH_2_‒) of MCC8400 showed a higher chemical shift (more deshielded) than MCC8399 (‒N‒CH_2_‒) [45]. Compound MCC7270 had the lowest E’ among the series of nitroimidazopyrazinones (−1662 mV). However, there was no direct correlation between the whole cell antitubercular activity and the ease of reduction of nitroimidazoles in our case, suggesting that factors other than redox activation are more crucial in determining their biological potential (Fig 3a).

**Fig 2.**
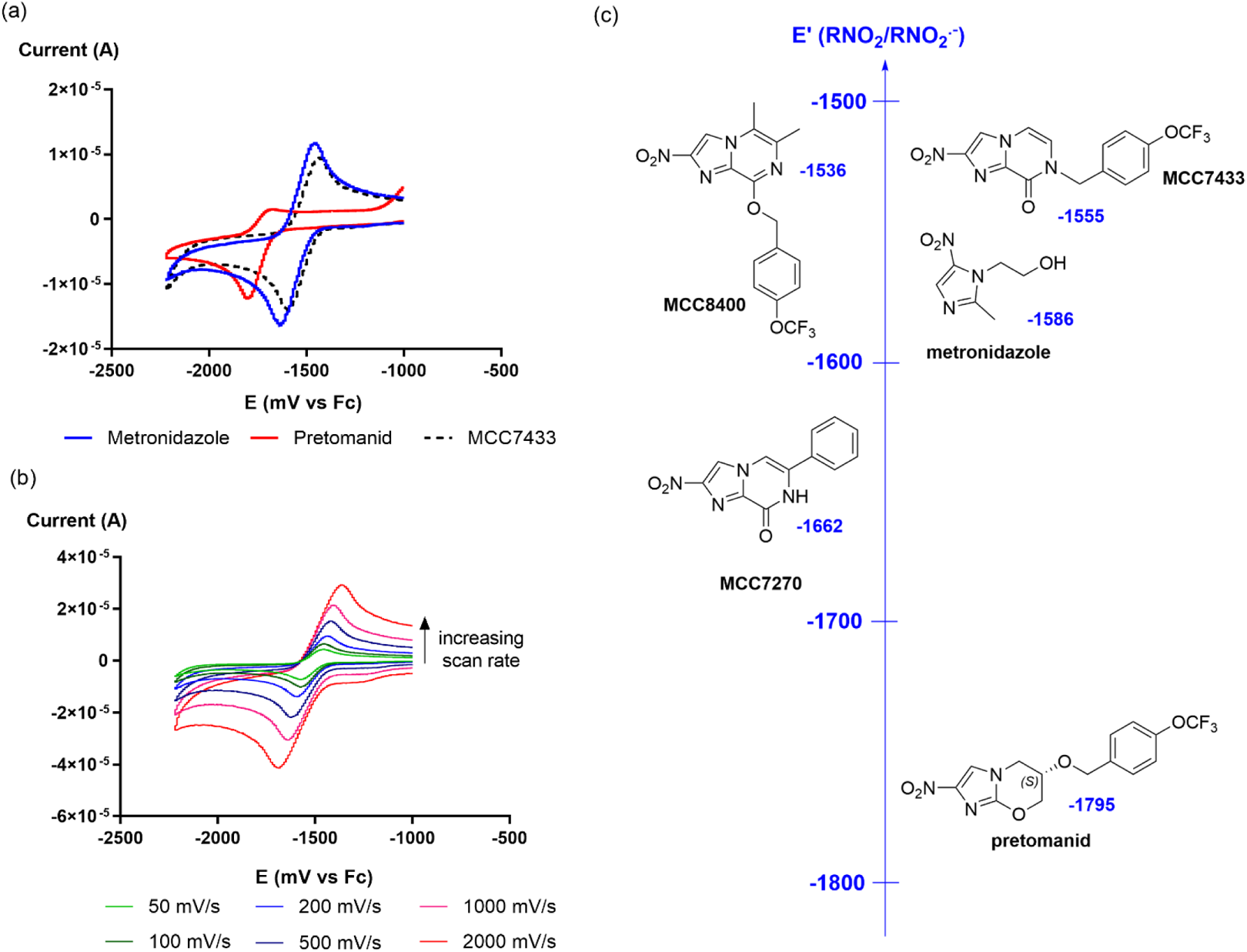
(a) Cyclic voltammogram of 1 mM tested compounds (metronidazole, pretomanid and MCC7433) in DMSO containing 0.1M tetrabutylammonium hexafluorophosphate (TBAHFP), with sweep rate at 200 mV/s. (b) Cyclic voltammogram of MCC7433 at different sweep rates (50, 100, 200, 500, 1000 and 2000 mV/s). (c) Relationship between electrochemical properties and structural variations of different nitroimidazoles.

**Fig 3.**
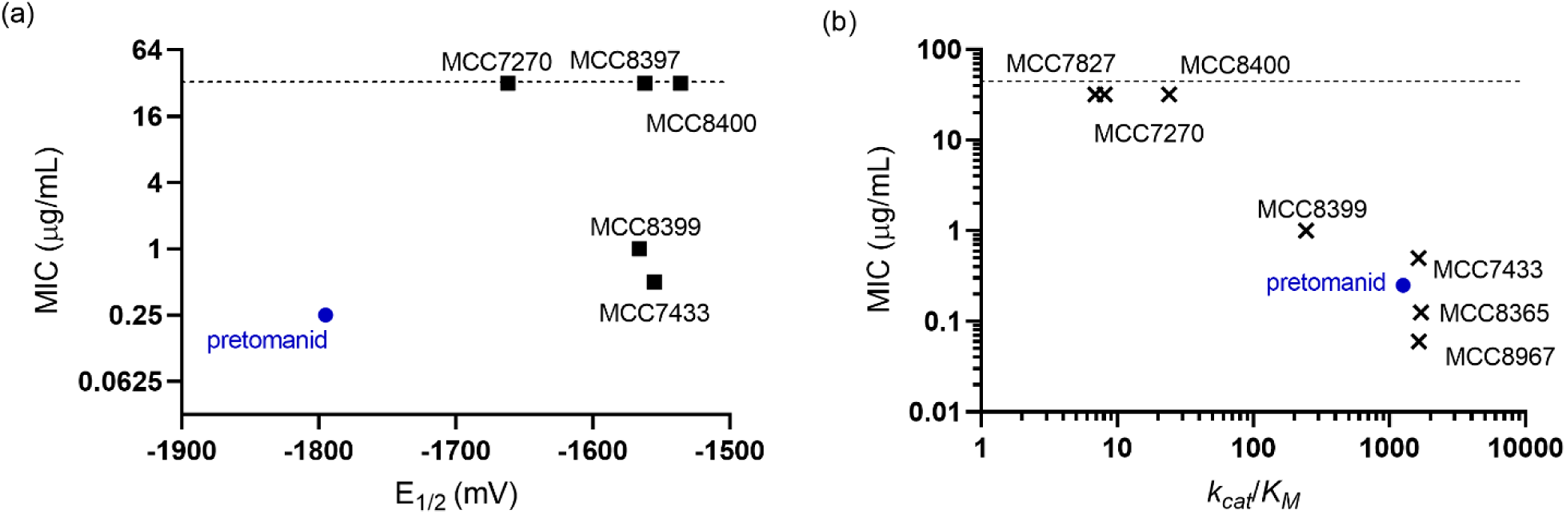
Correlation between the whole cell antitubercular activity (expressed as a log scale) and (a) redox potential (r = −0.291, R^2^ = 0.085, *p* = 0.634); (b) catalytic efficiency as Ddn substrates (r = −0.828, R^2^ = 0.686, *p* = 0.011). Dashed line represents the highest concentration tested (32 µg/mL) in minimum inhibitory concentration (MIC) assay.

**Table 2.** Characterization of Ddn activity and electrochemical properties of nitroimidazopyrazin-ones/-es.

| Compound | Core | R<sup>1</sup> | R<sup>2</sup> | R<sup>3</sup> | *k*<sub>cat</sub> (s<sup>-1</sup>) | *K*<sub>M</sub> (μM) | *k*<sub>cat</sub>/*K*<sub>M</sub> (M<sup>-1</sup> s<sup>-1</sup>) | E' (mV) vs Fc<sup>e</sup> |
| --- | --- | --- | --- | --- | --- | --- | --- | --- |
| Metronidazole |  |  |  |  | ns* | ns* | ns* | -1586 |
| Pretomanid | - | - | - | - | 9.3×10<sup>-3</sup> ± 4.8×10<sup>-4</sup> | 7.3 ± 1.4 | 1260 | -1795 |
| MCC7827 | - | - | - | - | 2.1×10<sup>-4</sup> ± 1.0×10<sup>-5</sup> | 31 ± 4.9 | 6.9 | ND |
| MCC7270 | A | H | Ph | H | 1.3×10<sup>-3</sup> ± 9.8×10<sup>-5</sup> | 164 ± 36 | 8.1 | -1662 |
| MCC7433 |  | 4-OCF<sub>3</sub>-benzyl | H | H | 3.3×10<sup>-3</sup> ± 1.4×10<sup>-4</sup> | 2.0 ± 0.41 | 1640 | -1555 |
| MCC8365 |  | 4-CH<sub>3</sub>-benzyl |  |  | 4.3×10<sup>-3</sup> ± 2.9×10<sup>-4</sup> | 2.5 ± 0.52 | 1710 | ND |
| MCC8364 |  | 3-OCF<sub>3</sub>-benzyl |  |  | 3.1×10<sup>-3</sup> ± 2.3×10<sup>-4</sup> | 45 ± 13 | 70 | ND |
| MCC8967 |  | 3-CF<sub>3</sub>-benzyl |  |  | 1.6×10<sup>-2</sup> ± 6.5×10<sup>-4</sup> | 9.4 ± 1.3 | 1650 | ND |
| MCC8397 |  | 4-OCF<sub>3</sub>-benzyl | CH<sub>3</sub> | H | ND | ND | ND | -1562 |
| MCC8399 |  | 4-OCF<sub>3</sub>-benzyl | CH<sub>3</sub> | CH<sub>3</sub> | 6.9×10<sup>-3</sup> ± 7.2 ×10<sup>-4</sup> | 28 ± 8.1 | 244 | -1566 |
| MCC8400 | B | 4-OCF<sub>3</sub>-benzyl |  |  | 7.0×10<sup>-4</sup> ± 6.4×10<sup>-5</sup> | 30 ± 6.8 | 24 | -1536 |

ns – not substrate; \*results obtained from literature [47]; ND – not determined.

As bioreduction of nitroimidazoles happens in an aqueous environment, the redox potential was further examined in a mixed medium, DMSO/PBS (70/30, v/v) with 0.1 M KCl as supporting electrolyte. However, the electrochemical reduction was irreversible due to the instability of radical anions in water in the presence of protons (data not shown). Multi-electron and proton associated irreversible reduction of nitro groups to give nitroso and hydroxylamine derivatives in aqueous solution is well established [46]. The lack of any reoxidation currents made reliable estimations of the redox potentials in aqueous solution problematic.

### Ddn enzymatic activity

It is known that the F_420_-dependent nitroreductase Ddn is responsible for the bioactivation of delamanid and pretomanid in *M. tuberculosis* [27, 28]. To investigate whether Ddn takes part in the activation of nitroimidazopyazinones, we first examined the efficiency of the most promising hit, MCC8967 as a substrate of this enzyme. We determined the kinetic activity of Ddn reducing pretomanid by following the oxidation of reduced F_420_, F_420_H_2._ We observed a *K_M_* for pretomanid of 7.3 μM, a *k*_cat_ of 9.3×10^−3^ s^−1^ and *k*_cat_*/K*_M_ of 1260 M^−1^s^−1^. This is similar to previously reported kinetic parameters of Ddn with pretomanid [47]. MCC8967 displayed a *K*_M_ of 9.4 μM indicating favorable binding with Ddn with a *k*_cat_ of 1.6×10^−2^. Its *k*_cat_*/K*_M_ of 1650 M^−1^ s^−1^ suggests that it is possibly a better substrate than pretomanid. To better understand the relationship between chemical structure and substrate efficiency, the kinetic parameters of structurally related analogs were explored (Table 2). The monocyclic nitroimidazole carboxamide MCC7827 was a poor substrate for Ddn, consistent with the inability of Ddn to oxidize F_420_H_2_ when metronidazole was used as substrate [47]. MCC7433 and MCC8365 were similar to MCC8967, with a *k_cat_/K_M_*of 1640–1710 M^−1^ s^−1^. However, when the trifluoromethoxy substituent was changed from a *para-*(MCC7433) to *meta-*(MCC8364) position, the binding affinity was reduced significantly from 2 μM to 45 μM, though antimycobacterial activity generally increased. The pyrazine analog MCC8400 and its pyrazinone counterpart MCC8399 shared similar *K_M_* values (30 μM and 28 μM respectively), although the *k*_cat_ of MCC8399 (6.9×10^−3^ s^−1^) was ten-fold higher than MCC8400 (7.0×10^−4^ s^−1^). There was no correlation between Ddn substrate specificity and the ease of reduction as determined from CV. However, in most of the cases (except MCC8364), their activity as substrates was quite closely associated with the whole cell potency against *M. tuberculosis* (Pearson’s r coefficient, r = −0.828, *p* = 0.011), indicating the potential importance of the Ddn enzyme in their activation (Fig 3b).

Previous studies have suggested that the extended side chains of pretomanid are favorable to accommodate the binding pocket of Ddn which is rather linear and elongated [48–50]. Therefore, a second generation of nitroimidazopyrazinones was generated to explore the role of the biaryl side chain of the nitroimidazopyrazinones in binding to the hydrophobic pocket of Ddn [35]. We tested selected analogs for their kinetic activity with Ddn and the results are summarized in Table 3. The biphenyl compound MCC9379 was found to have a stronger affinity (8.4 µM) and higher *k*_cat_/*K*_M_ (370 M^−1^ s^−1^) than the monoaryl MCC8364 (45 µM, 70 M^−1^ s^−1^), but this did not translate into having a lower MIC against *M. tuberculosis* (2 µg/ml and 0.125 µg/ml respectively). Replacement of one of the phenyl groups with a less lipophilic pyridine (MCC9380) or piperidine ring (MCC9347) resulted in marginal effects on both substrate specificity and cellular potency (5 µM and 4 µg/ml, 2.2 µM and 1-2 µg/ml respectively), with ~1–2-fold activity difference. Disrupting the molecular linearity and planarity of the extended side chains significantly improved the *k*_cat_/*K*_m_ value, with the 3’-linked phenylpyridine (MCC9426) showing 12-fold better activity (3190 M^−1^ s^−1^) compared to its linear 4’-linked counterpart MCC9380 (266 M^−1^ s^−1^). It is postulated that this orientation may be preferred over the linear side chains for favorable interactions with the active site of Ddn. Compound MCC9425 was the best substrate for Ddn in the present study, with a *k*_cat_/*K*_m_ of 4470 M^−1^ s^−1^, but was not the most potent at inhibiting *M. tuberculosis* growth (MIC 0.25 µg/ml).

**Table 3.** Characterization of nitroimidazopyrazinones with extended side chains.

| Compd | Form | X | link | R | $k_{\text{cat}}$ (s<sup>-1</sup>) | $K_{\text{M}}$ (μM) | $k_{\text{cat}}/K_{\text{M}}$<br>(M<sup>-1</sup> s<sup>-1</sup>) | Antibacterial<br>MIC (μg/mL) | |
| --- | --- | --- | --- | --- | --- | --- | --- | --- | --- |
|  |  |  |  |  |  |  |  | H37Rv<br>normoxia | H37Rv<br>hypoxia |
| **MCC9379** | A2 | C | 4' | 3-OCF<sub>3</sub> | $3.1 \times 10^{-3} \pm 3.1 \times 10^{-4}$ | $8.4 \pm 2.4$ | 370 | 2 (1-8) | 32 |
| **MCC9380** | A2 | N | 4' | 3-OCF<sub>3</sub> | $1.3 \times 10^{-3} \pm 7.2 \times 10^{-5}$ | $5.0 \pm 0.9$ | 266 | 4 (2-16) | >32 |
| **MCC9425** | A2 | N | 3' | 4-OCF<sub>3</sub> | $1.2 \times 10^{-2} \pm 4.1 \times 10^{-4}$ | $2.6 \pm 0.36$ | 4470 | 0.25 (0.25-1) | 82-88% I at 4-32 μg/mL |
| **MCC9426** | A2 | N | 3' | 3-OCF<sub>3</sub> | $1.4 \times 10^{-2} \pm 6.3 \times 10^{-4}$ | $4.3 \pm 0.57$ | 3190 | 2 (0.125-4) | 84% I at 32 μg/mL |
| **MCC9347** | A3 | - | - | - | $1.7 \times 10^{-3} \pm 7.8 \times 10^{-5}$ | $2.2 \pm 0.46$ | 773 | 1-2 | 8 |

% I indicates percentage of inhibition.

### Pharmacokinetic profile of MCC7270, MCC7433, MCC8364 and MCC8967

To further assess the potential for potent nitroimidazopyrazinones to be pursued as lead compounds for preclinical development, mouse *in vivo* pharmacokinetic (PK) studies were performed on analogs MCC7270, MCC7433, MCC8364 and MCC8967. These compounds have been previously evaluated for their drug-like properties [32] including microsomal stability, plasma stability, and plasma protein binding (Table 4). To ensure compound solubility prior to the PK study, MCC7433, MCC8364 and MCC8967 were formulated at 5 mg/mL in 10% DMSO and 90% PEG400, while MCC7270, which possessed higher aqueous solubility (Table 4), was prepared in a solution of 10% DMSO/40% PEG400/20% cremophor in water. The solutions were then administered to male CD-1 mice both orally and intravenously at 20 mg/kg and 5 mg/kg, respectively. Plasma samples were collected at different time points and analyzed by LC-MS/MS. The mean plasma concentration–time profiles are shown in Fig 4 and the key PK properties are summarized in Table 5. All compounds tested showed moderate to excellent oral bioavailability (F), with F = 64–98%. At a comparable dose, the more soluble MCC7270 reached the highest peak plasma concentration (C_max_) at 13 µg/mL and exposure (AUC_0-last_) at 88.3 µg.h/mL. MCC7270 also displayed similar absorption as pretomanid at T_max_ of 2 h (data obtained from literature [51]). When administered intravenously, the volume of distribution (V_ss_) of MCC7270 was 2.2-fold less than the total body water content (0.73 L/kg), indicating that it was more confined within the plasma, thus less distributed in tissues. Potent antitubercular analogs (MCC7433, MCC8364 and MCC8967) showed moderate to high absorption with a T_max_ of 2.7–5.3 h. Their oral C_max_ was in the range of 2.9‒3.4 µg/mL and exposure was within 15.6–38.9 µg.h/mL. Although the systemic plasma exposure of MCC7433 and MCC8967 was lower than MCC7270, their plasma concentrations remained above MIC values (ignoring any plasma binding effects) for at least 8 h (Fig 4). Therefore, they have the potential to be administered once daily.

**Fig 4.**
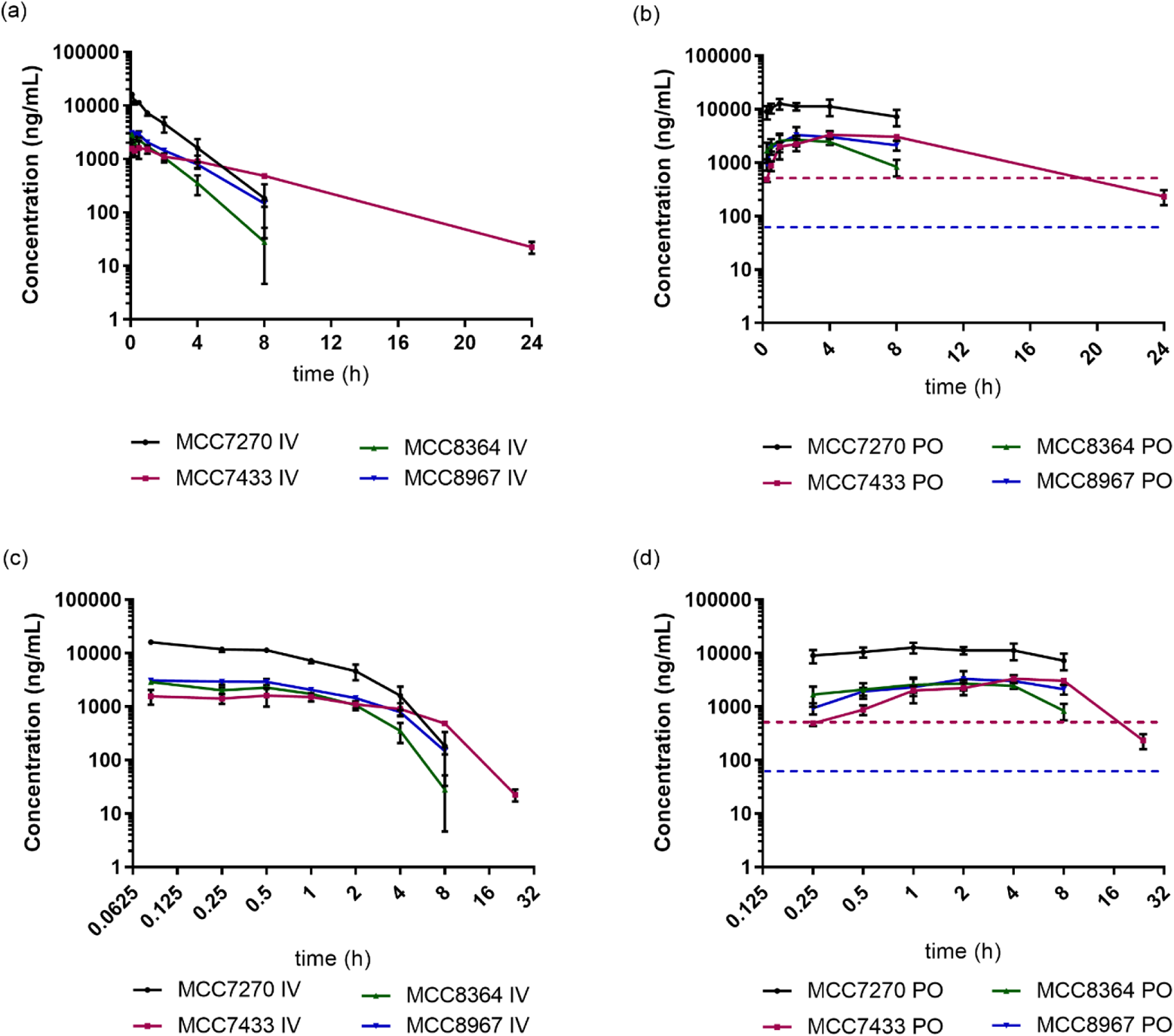
Plasma PK profiles of MCC7270, MCC7433, MCC8364 and MCC8967 after intravenous (IV, 5 mg/kg) and oral (PO, 20 mg/kg) administration in linear (a, b) and log (c, d) time scale. Data are shown as mean ± SD from three mice at different time points. Concentrations at 24 h were not determined for MCC7270, MCC8364 and MCC8967 as they were below the limit of quantitation. The in vitro MIC of MCC7433 and MCC8967 against M. tuberculosis H37Rv are shown as purple and blue dashed lines, respectively.

**Table 4.** *In vitro* ADME properties of pretomanid, MCC7270, MCC7433, MCC8364 and MCC8967.

| Compound | Solubility<br>($\mu\text{M}$) | PPB (%) | | Microsomal stability<br>(% remaining at 2 h) | | Plasma stability<br>(% remaining at 2 h) | |
| --- | --- | --- | --- | --- | --- | --- | --- |
|  |  | Human | Mouse | Human | Mouse | Human | Mouse |
| Pretomanid | 12 | 97 | 91 | 97 | 92 | 96 | 96 |
| MCC7270 | 27 | ND | ND | ND | ND | ND | ND |
| MCC7433 | <1 | >99 | >99 | >99 | >99 | 98 | >99 |
| MCC8364 | 5.4 | >99 | >99 | >99 | 97 | 99 | 97 |
| MCC8967 | 2.9 | 96 | 92 | ND | ND | ND | ND |

Values are presented as mean of three replicates. Some of the data were reported previously [32]. Solubility in water was determined by LC-UV (254 nm). Microsomal stability control, verapamil at 30 min = 9% (human), 2% (mouse). Plasma stability control, eucatropine at 2 h = 28% (human), 29% (mouse). % bound of PPB control, sulfamethoxazole = 57% (human), 9% (mouse). ND - not determined.

**Table 5.**
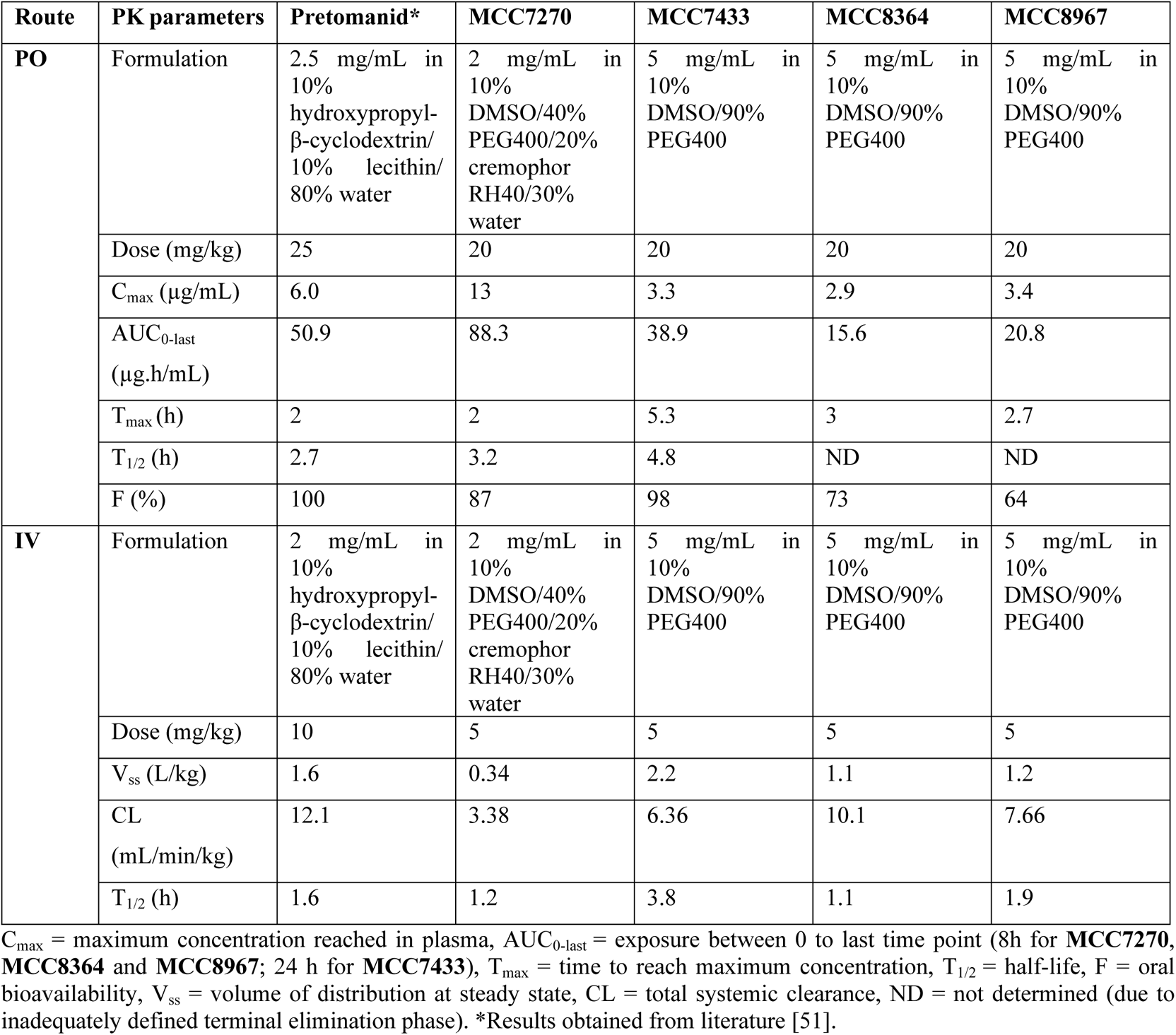
*In vivo* pharmacokinetic properties of selected analogs.

### Efficacy in TB mouse models

Two analogs, MCC7433 and MCC8967, were selected for assessment of efficacy in an acute BALB/c mouse *M. tuberculosis* infection model. These compounds were chosen based on their promising *in vitro* activity against both replicating and non-replicating *M. tuberculosis*, good PK profile as well as high *k*_cat_*/K*_M_ values with Ddn. *M. tuberculosis* was inoculated into the lung by aerosol at Day 0, and treatment started at Day 10, with pretomanid and rifampicin used as positive controls. Following three weeks of oral treatment, the levels of colony-forming units (CFU) in lungs were measured, with the test compounds at three different concentrations producing a dose-dependent response (Fig 5). The lung bacterial burden of the untreated group increased from 8.8 x 10^3^ CFU to 1.1 x 10^6^ CFU. MCC8967 was more efficacious than MCC7433 (*P* < 0.001), reducing the bacterial load by 0.75, 1.1 and 1.7 log CFU at 12.5, 25 and 50 mg/kg, respectively. At the highest dosing (50 mg/kg), MCC8967 showed comparable activity to pretomanid dosed at 20 mg/kg. MCC7433 was moderately active, with efficacy ranging from 0.21–0.72 log reduction in CFU compared to the vehicle control. Rifampicin was used as an additional control (Supporting information, Fig S1) and gave 2.7 log CFU reduction of bacterial load at 15 mg/kg. No mortality or obvious weight loss was found during the course of treatment, indicating the compounds were well-tolerated at the doses evaluated (Supporting information, Table S1).

**Fig 5.**
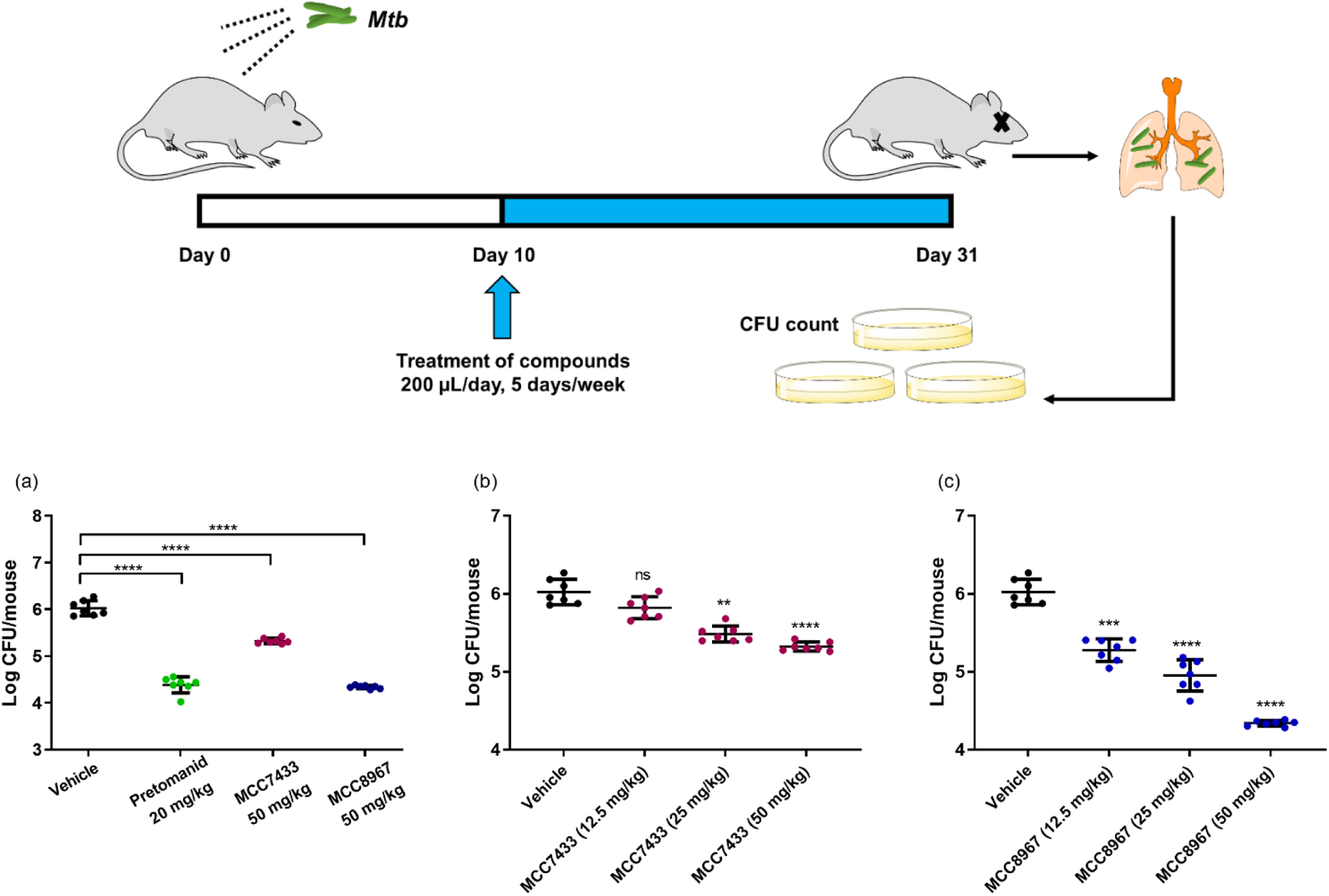
Treatment schedule in BALB/c mouse models of acute *M. tuberculosis* infection. *M. tuberculosis* was inoculated into the lung by aerosol at Day 0, and treatment started at Day 10. At day 31, mice were sacrificed and lungs were removed, homogenized, and plated onto 7H11 agar to determine the bacterial load. (a) CFU count data in the lungs of infected mice treated with vehicle, pretomanid, MCC7433 and MCC8967; dose response of (b) MCC7433 and (c) MCC8967 at 12.5, 25 and 50 mg/kg. Statistical significance was determined by ordinary one-way ANOVA followed by Dunnett’s multiple comparison test. \*\*\*\**P* < 0.0001, \*\*\**P* < 0.001, \*\**P* < 0.01, ns - not significant.

### Antitrypanosomal activity of extended side chains of nitroimidazopyrazinones

The repurposing potential of delamanid and pretomanid analogs for the treatment of kinetoplastid diseases [18, 20, 21] led us to investigate the antitrypansomal activity of nitroimidazopyrazinones against *T. cruzi*. Initial investigations using high content imaging demonstrated that analogs with extended biaryl side chains were highly active against *T. cruzi* (IC_50_ = 0.016–4.2 µM) [35]. This assay utilized an incubation time of 48 h, as longer incubation periods could bias the assay to identify *T*. *cruzi* cytochrome P450 (TcCYP51) inhibitors that are slow acting [52]. Early prediction of potential CYP inhibitors that are sub-efficacious is also possible with a 48 hour incubation [53]. As CYP51 is a promiscuous target in Chagas disease, targeting it has resulted in the failures of many clinical candidates including azole compounds (such as posaconazole). Therefore compounds that are fast-acting and long-lasting are often prioritised in drug discovery campaigns [54]. Of a total of 16 biaryl analogs, MCC9481 and MCC9482 were selected for further *in vivo* efficacy. MCC9481 was the most potent analog across the series when tested *in vitro* (IC_50_ of 0.016 µM, selectivity index of >2220), while MCC9482 (IC_50_ of 0.10 µM, selectivity index of >353) was favored due to its better solubility, especially in acidic conditions (>200 µM). These compounds completely cleared parasites (100%) from 3T3 host cells at E_max_ concentrations (maximal activity achievable in dose-response curves, Fig 6), in contrast to the CYP51 inhibitor, posaconazole that showed 87% inhibition at E_max_. This phenotypic characteristic suggested that MCC9481 and MCC9482 might not inhibit the CYP enzyme, consistent with other bicyclic nitroimidazole analogs that were reported to be inactive against CYP51 [21].

**Fig 6.**
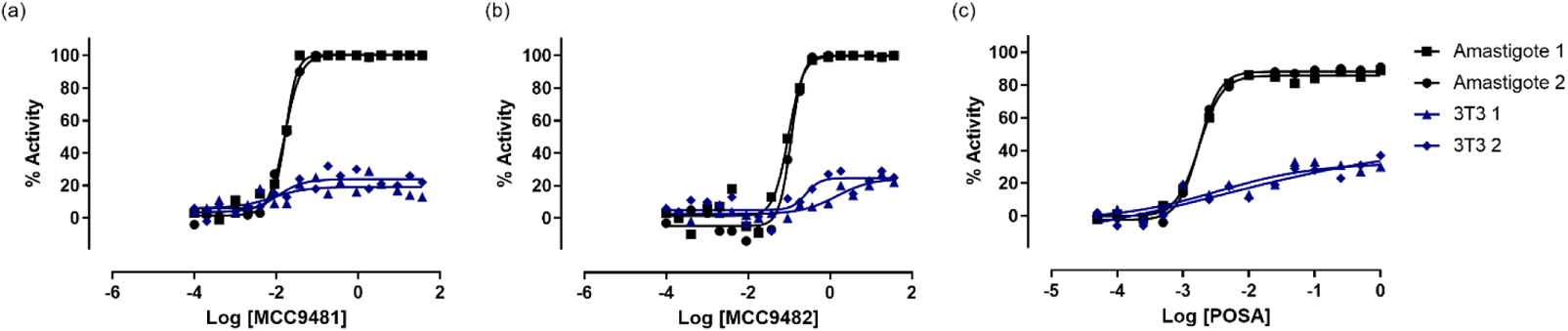
Dose-response curves for (a) MCC9481, (b) MCC9482, and (c) posaconazole (POSA) against *T. cruzi* amastigotes in 3T3 cells and their activity against 3T3 cells. The IC_50_ values of MCC9481 and MCC9482 were 0.016 ± 0.0003 µM and 0.10 ± 0.005 µM, with E_max_ of 100%.

### Pharmacokinetic profile of MCC9481 and MCC9482

The antiparasitic agents, MCC9481 and MCC9482 had distinctly different pharmacokinetic profiles (Fig 7, Table 6). MCC9482 had the fastest absorption rate (T_max_ = 0.67 h) among the nitroimidazopyrazinones investigated, while absorption of MCC9481 was ~10-fold slower, presumably due to its higher lipophilicity. Consistent with the mouse microsomal stability study (Table 7) [35], MCC9482 exhibited a short half-life (<1 h) when dosed orally. Despite having lower exposure due to its high clearance and high V_ss_, MCC9482 demonstrated good oral bioavailability at 92%. In contrast, MCC9481 had the longest half-life (6.1 h oral, 7.2 h IV) compared to other analogs including the monocyclic side chains MCC7270, MCC7374, MCC8364 and MCC8967. Plasma concentrations of MCC9481 reached a C_max_ at 5.3 h and remained above its *in vitro* IC_50_ for more than 24 h (Fig 7). However MCC9481 showed lower bioavailability (F = 45%), which is likely associated with its slower absorption and poor solubility.

**Fig 7.**
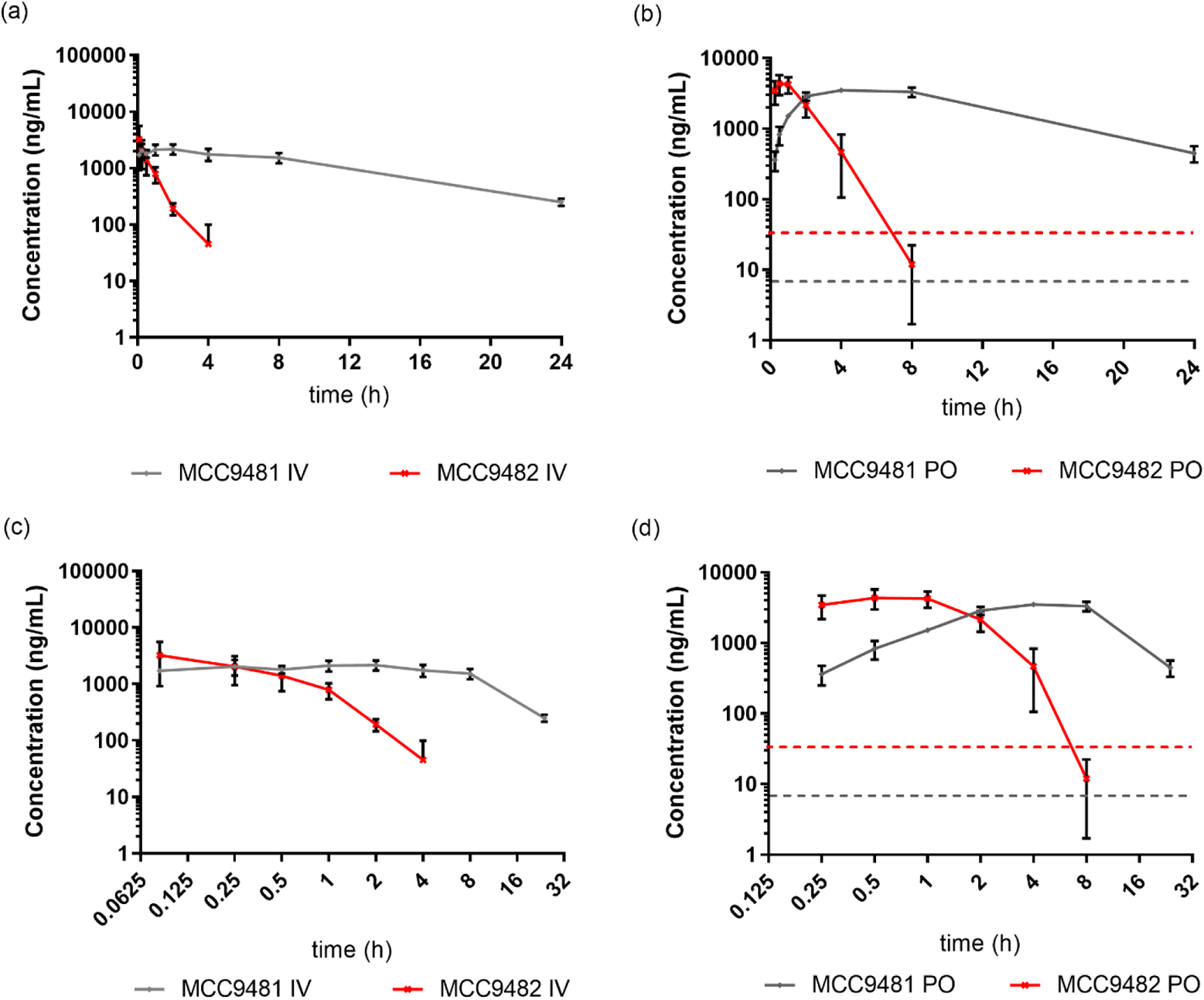
Plasma PK profiles of MCC9481 and MCC9482 after intravenous (IV, 5 mg/kg) and oral (PO, 20 mg/kg) administration in linear (a, b) and log (c, d) time scale. Data are shown as mean ± SD from three mice at different time points. Concentration at 24 h was not determined for MCC9482 as it is below the limit of quantitation. The *in vitro* IC_50_ of MCC9481 and MCC9482 against *T. cruzi* are shown as grey and red dashed lines.

**Table 6.** *In vivo* pharmacokinetic properties of anti-*T. cruzi* agents, MCC9481 and MCC9482.

| Route | PK parameters | MCC9481 | MCC9482 |
| --- | --- | --- | --- |
| PO | Formulation | 5 mg/mL in 10% DMSO/90% PEG400 | 5 mg/mL in 10% DMSO/90% PEG400 |
|  | Dose (mg/kg) | 20 | 20 |
| | $C_{max}$ (μg/mL) | 3.6 | 4.5 |
|  | AUC<sub>0-last</sub> (μg.h/mL) | 45.9 | 9.26 |
|  | T<sub>max</sub> (h) | 5.3 | 0.67 |
|  | T<sub>1/2</sub> (h) | 6.1 | 0.77 |
|  | F (%) | 45 | 92 |
| **IV** | Formulation | 5 mg/mL in 10% DMSO/90% PEG400 | 5 mg/mL in 10% DMSO/90% PEG400 |
|  | Dose (mg/kg) | 5 | 5 |
|  | V<sub>ss</sub> (L/kg) | 1.3 | 2.6 |
|  | CL (mL/min/kg) | 2.06 | 22.9 |
|  | T<sub>1/2</sub> (h) | 7.2 | 1.5 |

**Table 7.**
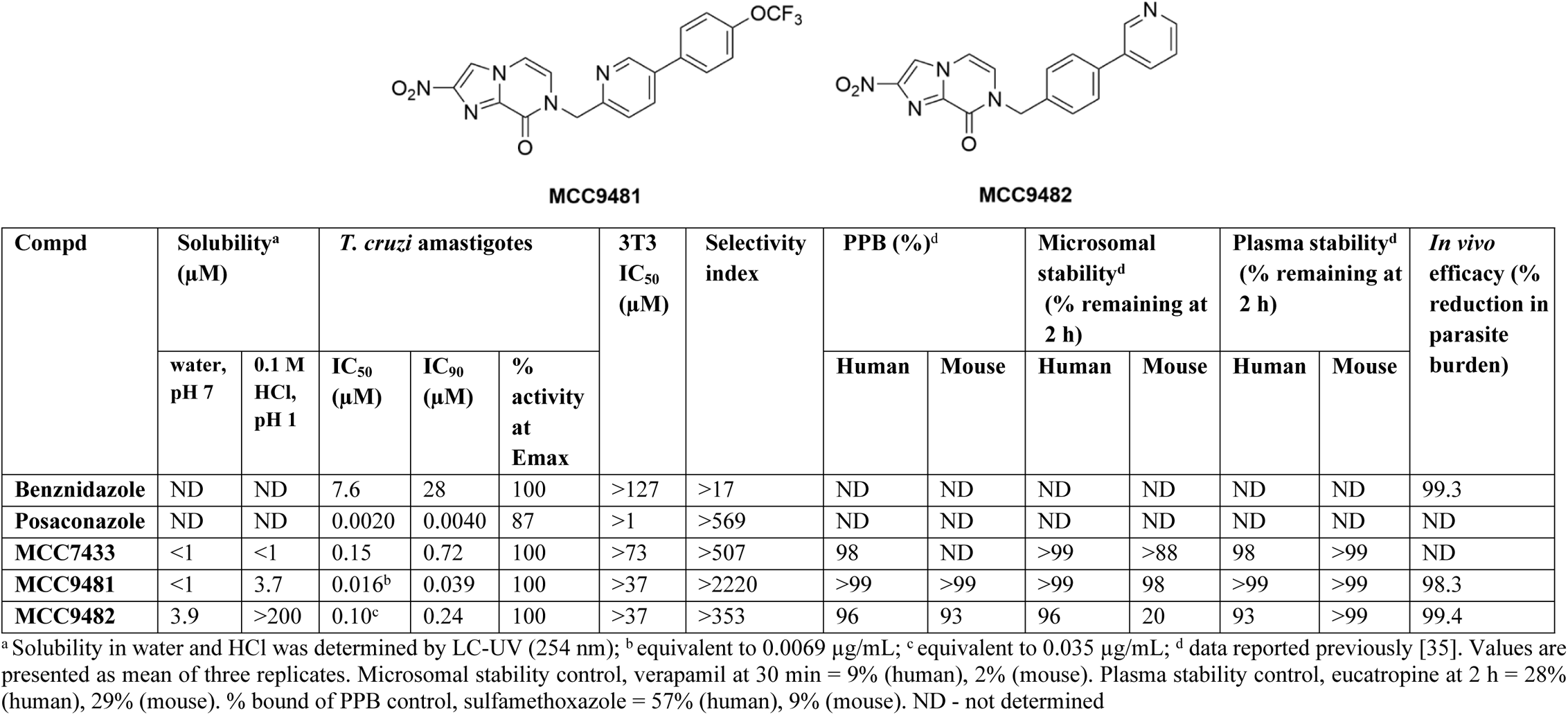
Activity profile of nitroimidazopyrazinones with biaryl side chains.

| Compd | Solubility<sup>a</sup><br>(μM) |  | *T. cruzi* amastigotes |  |  | 3T3<br>IC<sub>50</sub><br>(μM) | Selectivity<br>index | PPB (%)<sup>d</sup> |  | Microsomal<br>stability<sup>d</sup><br>(% remaining at<br>2 h) |  | Plasma stability<sup>d</sup><br>(% remaining at<br>2 h) |  | *In vivo*<br>efficacy (%<br>reduction in<br>parasite<br>burden) |
| --- | --- | --- | --- | --- | --- | --- | --- | --- | --- | --- | --- | --- | --- | --- |
|  | water,<br>pH 7 | 0.1 M<br>HCl,<br>pH 1 | IC<sub>50</sub><br>(μM) | IC<sub>90</sub><br>(μM) | %<br>activity<br>at<br>Emax |  |  | Human | Mouse | Human | Mouse | Human | Mouse |  |
| **Benznidazole** | ND | ND | 7.6 | 28 | 100 | >127 | >17 | ND | ND | ND | ND | ND | ND | 99.3 |
| **Posaconazole** | ND | ND | 0.0020 | 0.0040 | 87 | >1 | >569 | ND | ND | ND | ND | ND | ND | ND |
| **MCC7433** | <1 | <1 | 0.15 | 0.72 | 100 | >73 | >507 | 98 | ND | >99 | >88 | 98 | >99 | ND |
| **MCC9481** | <1 | 3.7 | 0.016<sup>b</sup> | 0.039 | 100 | >37 | >2220 | >99 | >99 | >99 | 98 | >99 | >99 | 98.3 |
| **MCC9482** | 3.9 | >200 | 0.10<sup>c</sup> | 0.24 | 100 | >37 | >353 | 96 | 93 | 96 | 20 | 93 | >99 | 99.4 |

<sup>a</sup> Solubility in water and HCl was determined by LC-UV (254 nm); <sup>b</sup> equivalent to 0.0069 μg/mL; <sup>c</sup> equivalent to 0.035 μg/mL; <sup>d</sup> data reported previously [35]. Values are presented as mean of three replicates. Microsomal stability control, verapamil at 30 min = 9% (human), 2% (mouse). Plasma stability control, eucatropine at 2 h = 28% (human), 29% (mouse). % bound of PPB control, sulfamethoxazole = 57% (human), 9% (mouse). ND - not determined

### *In vivo* efficacy study in a bioluminescent *T. cruzi* mouse model

As a proof of concept, MCC9481 and MCC9482 were evaluated for their *in vivo* activity using an acute *T. cruzi* infected murine model and bioluminescence imaging. BALB/c mice (n=3/group) infected with bioluminescent *T. cruzi* CL Brener parasites (see Materials and Methods) were treated at the peak of the acute stage infection (14 dpi). The untreated group (vehicle only) displayed high parasitemia levels until 24 dpi, whereas mice treated with MCC9481 and MCC9482 showed a reduction in parasite burden of >98‒99% by 18 dpi that persisted until 24 dpi (Fig 8). A similar outcome was observed in mice treated with benznidazole at 100 mg/kg. However, all mice were bioluminescence positive by 36 dpi. Both MCC9481 and MCC9482 were able to transiently reduce the parasite burden to background levels by the end of treatment, with no significant adverse effects observed in all mice treated with both compounds (a slight weight loss was noticed in mice treated with benznidazole). Although not significant, at 36 dpi the bioluminescence intensity was more intense in the mice treated with benznidazole compared to the mice that were treated with both MCC9481 and MCC9482 (Fig 8).

**Fig 8.**
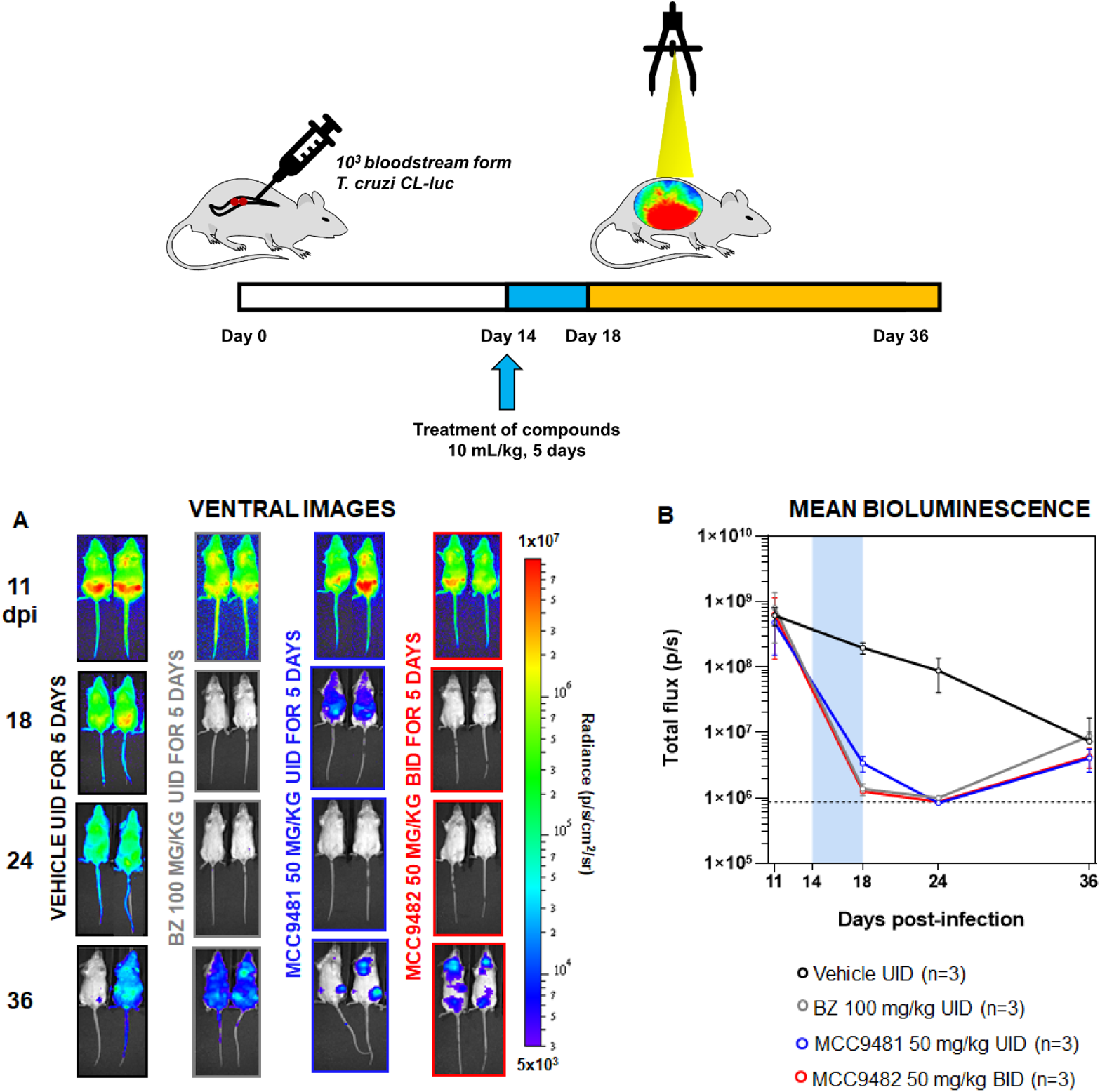
In vivo assessment of compounds MCC9481 and MCC9482 as a treatment for experimental acute stage *T. cruzi* infections. (A) Representative ventral images of two BALB/c mice infected with bioluminescent CL-Brener were treated at the acute stage of infection (day 14) with vehicle (orally, 5 days, once daily, n=3), benznidazole (BZ) at 100 mg/kg (orally, 5 days, once daily, n=3), MCC9481 at 50 mg/kg (orally, 5 days, once daily, n=3) and MCC9482 at 50 mg/kg (orally, 5 days, twice daily, n=3) and followed until 36 days post-infection (dpi). Heat-maps are on log^10^ scales and indicate intensity of bioluminescence from low (blue) to high (red). At day 36 post-infection, all mice were humanely killed. (B) Mean bioluminescence evaluation of mice treated with MCC9481 and MCC9482 during acute stage *T. cruzi* infections. Treatment groups, dose regimens, including time of treatment (blue bar), are indicated.

## Conclusion

Nitroimidazoles were once considered to be ‘undesirable’ in drug discovery due to the possible toxicity issues associated with the nitro group. However, recent advances have demonstrated that new nitroaromatics with favorable therapeutic indices are possible by controlling the bioactivation of the nitro groups. This study has expanded our prior work to further explore the properties of nitroimidazopyrazinones as a new bicyclic subclass capable of treating both tuberculosis and CD. Nitroimidazopyrazinones were found to be highly selective at killing *M. tuberculosis*, but were not active against nontuberculosis mycobacteria. Through enzymatic screening against deazaflavin-dependent nitroreductase (Ddn), a subset of nitroimidazopyrazinones were identified as effective substrates. Although there was some good correlation between enzymatic activity and mycobacteria killing, it should be noted that the kinetic parameters of Ddn is only a reflective of F_420_H_2_ reoxidation and does not translate directly into the whole cell activity. A more detailed biochemical analysis such as measuring the kinetics of NO production by Ddn may provide greater insight into their mode of action in *M. tuberculosis*. In the future, a comparison of activity against Ddn mutants or other homologues such as Rv1261c, Rv1558 and Rv3178 could assess the specificity of this Ddn-mediated bioactivation. As resistance towards nitroimidazoles have been found to already be present due to mutations in Ddn, assessing and developing new compounds that can be activated by more than one F_420_-dependent enzyme would be of great benefit [39, 55].

Electrochemical properties were determined by cyclic voltammetry but there was not a clear correlation between ease of reduction of nitroimidazoles and their bioactivity. It is possible that nitroimidazopyrazinones exert their activity through several pathways, and therefore a single parameter (such as Ddn catalytic efficiency and redox potential) will not correlate perfectly to their potency. These multiple targeting mechanisms can potentially be used to overcome or delay the rise of resistance.

Compound MCC8967 was identified as the most promising hit compound against *M. tuberculosis*, with satisfactory pharmacokinetic profile and high efficacy in acute TB mouse models. A different subset of nitroimidazopyrazinones, with extended biaryl side chains, demonstrated sub-micromolar inhibitory activity against *T. cruzi in vitro* and were able to suppress infection *in vivo* in a mouse model. MCC9482, with a terminal pyridine side chain (pyridine-3-yl-phenyl group), demonstrated improved solubility and oral bioavailability (F=92%) compared to MCC9481 (4-trifluromethoxyphenyl-3-pyridine group), while MCC9481 displayed a longer half-life and could be used for once daily dosing. Due to the absence of Ddn in *T. cruzi*, it was postulated that other nitroreductases such as TcNTR-1 [56], might involve in the bioactivation of these bicyclic nitroimidazoles.

Taken together, our work has revealed the potential of a new bicyclic nitroimidazole subclass for the treatment of tuberculosis and Chagas disease, supported by *in vivo* pharmacokinetic and efficacy studies that demonstrate the potential for active nitroimidazopyrazinones to be developed as clinical candidates. Further exploration of their multiple mode of actions under both aerobic and hypoxic conditions could help understand their capacity to target drug-resistant organisms, while additional *in vivo* studies could optimise dosing regimens and evaluate efficacy against both chronic and acute infections.

## Materials and Methods

### Synthesis of nitroimidazopyrazinones

Nitroimidazopyrazinone analogs were synthesized according to our recently reported methods [35, 45]. The purity of these compounds was >95% as determined by liquid chromatography-mass spectrometry (LC-MS) using UV at 254 nm, ELSD and APCI/ESI-MS detectors. Synthetic routes and experimental details are briefly described in Supporting Information Scheme S1.

### Mycobacterial minimum inhibitory concentration assay

The minimum inhibitory concentration (MIC) study was performed against *M. tuberculosis* H37Rv (ATCC 27294), avirulent *M. tuberculosis* H37Ra (ATCC 25177), *M. avium* (ATCC 25291) and *M. smegmatis* mc^2^155 (ATCC 700084) using a resazurin reduction microplate assay as previously described [32]. Compounds were serially diluted across the 96-well microtiter plates in 100 µL of 7H9S media (7H9 with 10% ADC, 0.5% glycerol, 0.05% Tween-80 and 1% tryptone). *M. tuberculosis* H37Rv, *M. tuberculosis* H37Ra, *M. avium* and *M. smegmatis* were was cultured in Middlebrook 7H9 broth medium supplemented with ADC (Difco Laboratories), 0.5% glycerol, and 0.02% Tyloxapol at 37 °C until mid-exponential phase (OD_600_ 0.4-0.8) before being diluted to the appropriate cell density in 7H9S media and added to the assay plate. Final cell density in the assay plates was ~2×10^4^ CFU/mL (OD_600_ 0.001) for *M. tuberculosis* H37Rv, H37Ra & *M. avium* and 4×10^4^ CFU/mL (OD_600_ 0.002) for *M. smegmatis.* The plates were then incubated for 5 days for *M. tuberculosis* H37Rv, 6 days for *M. tuberculosis* H37Ra and *M. avium*, and 24 h for *M. smegmatis* at 37 °C in a humidified incubator before the addition of 30 μL of a 0.02% resazurin solution and 12.5 μL of 20% Tween-80 and further incubated for an additional 24 h at 37 °C. Sample fluorescence was measured on a Fluorostar Omega fluorescent plate reader (BMG) (excitation 530 nm, emission 590 nm).

For hypoxic assays of *M. tuberculosis* H37Rv, the same method was used as the normoxic condition except the plates were incubated for 5 days at 0.1% oxygen before the addition of resazurin and incubated for an additional 48 h prior to fluorescence reading. Percent fluorescence relative to the positive control wells (bacteria without compound) minus the negative control wells (media only) was plotted for the determination of the MIC, the lowest concentration at which the percentage inhibition was ≥90%.

### Cyclic voltammetry analysis

The electrochemical study was done using a BAS CV-100 voltammetric analyser. All the voltametric measurements were performed at room temperature in a 5 mL voltametric cell. Oxygen was removed by bubbling argon gas through the sample solution before measurements were taken. Reference electrode used for experiments in anhydrous DMSO and mixed medium (mixture of 70% DMSO and 30% of PBS) was Ag/AgNO_3_ and Ag/AgCl, respectively. A glassy carbon electrode (BAS) was used as working electrode and a platinum wire served as counter electrode. BAS was polished with 0.05 µm alumina prior to and in the interval of experiments. Supporting electrolytes used were 0.1 M tetrabutylammonium hexafluorophosphate (TBAHFP) for measurements in DMSO and 0.1 M KCl for mixed medium. Compounds were tested at 1 mM concentration. Ferrocene was used as the reference point for the formal potential (E′) measurements in DMSO. Different sweep rates at 50-2000 mVs^−1^ were used.

### Protein expression and purification

Deazaflavin-dependent nitroreductase (Ddn) was expressed and purified as previously described [39]. In brief, Ddn enzyme was transformed into *Escherichia coli* BL21 (DE3) and grown on ampicillin-containing Luria-Bertani (LB) agar plates. Single colonies were picked, inoculated in LB media with ampicillin (100 µg/mL) and grown overnight. The overnight cultures were diluted 100x and grown until OD_600_ 0.4. Expression was induced with isopropyl β-D-1-thiogalactopyranoside (IPTG) to a final concentration of 0.3 mM. Cultures were grown for 3 h at 25 °C before harvested by centrifugation at 8,500 × *g* for 20 min at 4 °C. Cells were then suspended in lysis buffer and sonicated using an Omni Sonicator Ruptor 400 (2 x 6 min. at 50% power). The lysed cell suspensions were centrifuged at 13,500 × *g* for 1 h at 4 °C. Soluble extract was obtained from supernatant and purified using amylose resin (NEB). The purified protein was frozen at −80 °C in in 20 mM Tris (pH 7.5), 200 mM NaCl, 10 mM maltose, and 10% glycerol. Cofactor F_420_ was purified from M. *smegmatis* mc24517 following published protocols [57, 58]. F420–dependent glucose-6-phosphate dehydrogenase (FGD) was expressed and purified as described earlier [39].

### Nitroreductase enzymatic assay

The nitroreductase enzyme assay was performed by monitoring oxidation of F_420_H_2_ spectrophotometrically using fluorescence (Ex/Em 420/470 nm) as previously described [39, 59]. F_420_H_2_ was first prepared by overnight reduction of F_420_ with 10 µM FGD and 10 mM glucose-6-phosphate in 20 mM Tris-HCl (pH 7.5) under anaerobic conditions. FGD was removed by spin filtration in a 0.5-mL 10K molecular-weights cutoff (MWCO) spin filter (Millipore), and F420H2 was used within 1 h of FGD being removed. The reaction was initiated at room temperature by adding 0.1‒1 µM of Ddn enzyme to the assay mixture containing 25 µM of F_420_H_2_ and 0-300 µM of substrate in 200 mM Tris-HCl (pH 7.5), 0.1 % Triton X-100. Control reactions without Ddn and without substrate were also performed. Specific enzyme activity of F_420_H_2_ oxidation was calculated using a standard curve of F_420_. To determine the correlation between *k_cat_/K_m_* values and its whole cell potency (MIC), Pearson’s r-coefficient and statistical significance (*p*) were calculated using GraphPad Prism 9 software (San Diego, CA, USA).

### *In vivo* pharmacokinetic study

The mouse pharmacokinetic study was conducted by WuXi AppTec Co., Ltd. (Shanghai). The experimental procedures were approved by the Institutional Animal Care and Use Committee of WuXi AppTec Co., Ltd. (Protocol number PK01-001-2019v1.0). Compounds were administered to CD-1 male mice via oral (PO) and intravenous (IV) routes, in 20 mg/kg and 5 mg/kg respectively. Compounds MCC7433, MCC8364, MCC8967, MCC9481 and MCC9482 were formulated in 10% DMSO and 90% PEG400, whereas MCC7270 was in a solution of 10% DMSO/40% PEG400/20% cremophor in water. Plasma samples were taken at different time points (IV: 0.083, 0.25, 0.5, 1, 2, 4, 8 and 24 h; PO: 0.25, 0.5, 1, 2, 4, 8 and 24 h). Briefly, 30 μL of blood was taken via submandibular or saphenous vein for the first several time points whereas blood samples at the last time point (24 h) were collected via cardiac puncture when the mouse was under terminal anaesthesia. All blood samples were transferred into pre-chilled microcentrifuge tubes containing 2 μL of 0.5 M of K_2_-EDTA as anti-coagulant and were centrifuged at 7,000 rpm, 4 °C for 10 min. Plasma was then collected, frozen over dry ice, and stored at −70 °C until LC-MS/MS analysis. To process the samples prior to analysis, an aliquot of 5 µL was quenched with 300 µL of acetonitrile containing internal standards (labetalol, tolbutamide, verapamil, dexamethasone, glyburide and celecoxib, 100 ng/mL for each). The mixture was vortex-mixed and centrifuged for 15 min at 4,000 rpm, 4 °C. Supernatant (50 µL) was transferred to 96-well plate and centrifuged again for 5 min at 4,000 rpm, 4 °C before injecting to LC-MS/MS. Calibration curve was prepared for quantitation. Mobile phase was 0.1% formic acid & 2 mM ammonium formate in water/acetonitrile (v:v, 95:5) (solvent A) and 0.1% formic acid & 2 mM ammonium formate in acetonitrile/water (v:v, 95:5) (solvent B). LC-MS/MS condition: column ACQUITY UPLC HSS T3 1.8 μm 2.1 × 50 mm; column temperature: 60 °C; flow: 0.6 mL/min; gradient timetable: 0.00 min, 10% B; 1.20 min, 90% B; 1.40 min, 90% B; 1.41 min, 10% B; 1.50 min, 10 % B. Pharmacokinetic data were calculated using Phoenix WinNonlin 6.3 (Certara, Princeton, USA).

### *In vivo* acute TB mouse study

The experimental procedures for acute TB mouse efficacy study were approved by the Animal Care Policies of the University of Illinois at Chicago (ACC Number: 18-135). Female BALB/c mice weighing 21–23 g were infected with *M. tuberculosis* by aerosol. Compounds were formulated in 5% DMSO and 10% hydroxypropyl-β-cyclodextrin prior to administration. The *in vivo* efficacy study was performed according to a published method [60]. At day 10 post-infection, each compound was given orally in 200 µL at different doses (12.5, 25 and 50 mg/kg). The treatment was continued for 3 weeks, 5 days a week. Each treated group composed of 7 mice. Pretomanid was used as the comparator and was dosed at 20 mg/kg. Rifampicin (15 mg/kg) was used as a positive control. The mice were humanely killed after 31 days and the lungs were removed, homogenized and diluted in HBSS before plated onto 7H11 agar. Efficacy results are presented as colony forming unit (CFU) and was plotted using GraphPad Prism 7 (San Diego, CA, USA). Statistical analysis was evaluated by a one-way analysis of variance (ordinary one-way ANOVA), followed by a Dunnett’s multiple comparison test to compare the differences between untreated and experiment groups. Results were considered statistically significant at 95% confidence level.

### *T*. *cruzi* image-based assay

Compounds were screened against *T. cruzi* intracellular amastigotes using an established image-based assay as described previously [35, 53]. Briefly, 3T3 fibroblasts (ATCC CCL92) were infected with *T. cruzi* Tulahuen strain parasites at a multiplicity of 5:1. Compounds, pre diluted in milli-Q H2O, were added to wells to give final assay concentrations ranging in serial log dilutions from 73.3 or 36.6 µM, to 1.8×10-4 µM or 9×10-5 µM, respectively. Plates were incubated for 48 h before fixing infected cells with 4% paraformaldehyde containing Hoechst 3348, followed by staining with HCS CellMask™ green (ThermoFisher Scientific). Images were taken on a Phenix high-content cell imager (PerkinElmer). All experiments were performed over two biological replicates. IC_50_ values were calculated in GraphPad Prism 9 (San Diego, CA, USA), using a sigmoidal dose-response analysis, with variable slope.

### *In vivo* bioluminescence study of experimental acute stage *Trypanosoma cruzi* infections

All animal work was performed under UK Home Office license PPL P9AEE04E4 and approved by the London School of Hygiene and Tropical Medicine Animal Welfare and Ethical Review Board (AWERB). All protocols and procedures were conducted in accordance with the UK Animals (Scientific Procedures) Act 1986. The study was performed with a slight modification as previously described [61]. Female BALB/c mice (n=3/group) were purchased from Charles River (UK) and maintained under specific pathogen-free conditions in individually ventilated cages. They experienced a 12-hour light/dark cycle and had access to food and water *ad libitum*. Bioluminescent *T. cruzi* CL Brener parasites were generated by genetic transformation with the construct pTRIX2-RE9h [62]. Mice were injected i.p. with 10^3^ bloodstream form trypomastigotes (in 0.2 mL in D-PBS) obtained from infected SCID mice. MCC9481 and MCC9482 were formulated in 5% DMSO and 10% hydroxypropyl-β-cyclodextrin at 5 mg/mL and prepared daily before administration. Benznidazole was formulated in an aqueous suspension vehicle containing 0.5% (w/v) hydroxypropyl methylcellulose and 0.4% (v/v) Tween 80 at 10 mg/mL on the first day of treatment and left at 4 °C until needed. When infections had reached the acute stage at day 14 post-infection, each compound was administered by oral gavage (adjusted by weight), and vehicle only was administered to control mice (negative control), for 5 days at 50 mg/kg for MCC9481 and MCC9482 and 100 mg/kg for benznidazole (positive control). MCC9481 was given once a day (UID), while MCC9482 was given twice daily (BID) due to its lower half-life (Table 7). For bioluminescence imaging, mice were injected with 150 mg/kg d-luciferin i.p., then anaesthetized using 2.5% (v/v) gaseous isoflurane in oxygen for 2-3 min. Mice were placed in an IVIS Spectrum system (Caliper Life Science) and images acquired 10-20 min after d-luciferin administration using Living Image^®^ 4.3. Exposure times varied from 10 secs to 5min, depending on signal intensity. After imaging, mice were returned to cages. To estimate parasite burden, regions of interest (ROIs) were drawn using Living Image^®^ 4.3 to quantify bioluminescence expressed as total flux (photons/second; p/s). The detection threshold was established from uninfected mice.

## Acknowledgements

We are thankful to the support from the University of Queensland, Australian National University, University of Illinois at Chicago, London School of Hygiene & Tropical Medicine, and Griffith University for providing facilities. We also thank Angie Jarrad for helpful discussions at the early stage of this work.

## Supporting Information

Table S1. Average body weight of mice at day 10 (starting of treatment) and day 31 (end of treatment) for *M. tuberculosis* infection model.

Fig S1. CFU count data in the lungs of infected mice treated with rifampicin. Scheme S1. General reaction scheme of nitroimidazopyrazinones.

